# Isolation of strains and their genome sequencing to analyze the mating system of *Ophiocordyceps robertsii*

**DOI:** 10.1101/2022.12.02.518814

**Authors:** Melvin Xu, Nathan A. Ashley, Niloofar Vaghefi, Ian Wilkinson, Alexander Idnurm

## Abstract

The fungal genus *Ophiocordyceps* contains a number of insect pathogens. One of the best known of these is *Ophiocordyceps sinensis*, which is used in Chinese medicine and its overharvesting threatens sustainability; hence, alternative sources are being sought. *Ophiocordyceps robertsii*, found in Australia and New Zealand, has been proposed to be a close relative to *O. sinensis*, but little is known about this species despite being also of historical significance. Here, *O. robertsii* strains were isolated into culture and high coverage draft genome sequences obtained and analyzed. This species has a large genome expansion, as also occurred in *O. sinensis*. The mating type locus was characterized, indicating a heterothallic arrangement whereby each strain has an idiomorphic region of two (*MAT1-2-1, MAT1-2-2*) or three (*MAT1-1-1, MAT1-1-2, MAT1-1-3*) genes flanked by the conserved *APN2* and *SLA2* genes. These resources provide a new opportunity for understanding the evolution of the expanded genome in the homothallic species *O. sinensis*, as well as capabilities to explore the pharmaceutical potential in a species endemic to Australia and New Zealand.

**One sentence summary:** *Ophiocordyceps robertsii* is a close relative of *O. sinensis* and has a large genome but with a heterothallic mating system.

## INTRODUCTION

*Ophiocordyceps robertsii* is an entomopathogenic fungus of historical note and of potential pharmaceutical value. *O. robertsii* is pathogenic to ghost moth larvae in the Hepialidae family, is endemic in Australia and New Zealand, and was the first fungus described from New Zealand by Europeans [1]. When assigned its Linnaean name, the fungus was already well known to the Māori culture, who used *āwheto* as a source of ink for their ritual tattoos [2].

*O. robertsii* was proposed as the closest relative of *Ophiocordyceps sinensis* [3], which is known as *dōng chóng xià cǎo* (‘summer grass, winter worm’) and is China’s, if not the world’s, most valued medicinal fungus. On parity with the cost of gold, this is the most valuable edible fungal product by weight, well over that of white truffles [4]. With its limited geographic distribution on the Qinghai-Tibetan Plateau, *O. sinensis* harvesting has forced extensive pressures into this sensitive region and on this species. Producing this fungus in cultivation may protect its natural habitat [5, 6]. Alternatively, closely related species may also provide a new resource for use in medicine and/or as origins for new pharmaceutical chemicals.

The biology, population structure, diversity and relationships with other species of *O. robertsii* are unknown. In particular, the life cycle of *O. robertsii* is uncharacterized. If to be used as a source of beneficial chemicals, e.g. as anticancer agents [7], that would ideally be isolated from *in vitro* cultures it is essential to determine the mating system, especially as some fungal secondary metabolites may only be produced at certain stages of the life cycle. When comparing *O. robertsii* with *O. sinensis*, a number of questions emerge. The genome of *O. sinensis* shows an expansion in size relative to other members of the *Ophiocordycipitaceae* family and the species has a homothallic life style [8, 9]: how did this arise? Which species, as currently known, is the closest relative to *O. sinensis*, and could that species be informative for a better understanding of the evolution of species within the *Ophiocordyceps* genus? With these questions in mind, the primary aim of this research was to define the mating system of *O. robertsii*, by developing genomic resources that could also contribute further to understanding the evolution of entomopathogenic fungi.

## MATERIALS AND METHODS

### Fungal material

A fruiting body of *O. robertsii* and associated parasitized larva was collected in 2018 near Queenstown, Tasmania (under Tasmanian Department of Primary Industries, Parks, Water and Environment permit FL 18158). A second fruiting body was donated by a member of the public, having been collected in an undescribed location in southeastern Victoria (received under Victorian Department of Environment, Land, Water and Planning permit 10008557). The fruiting bodies were pressed onto potato dextrose agar (PDA) medium supplemented with chloramphenicol and cefotaxime (30 µg/ml and 100 µg/ml, respectively) to reduce bacterial growth, to release ascospores. Colonies derived from single ascospores were excised with a scalpel and transferred to new PDA plates.

### Insect identification

Genomic DNA was extracted from the posterior end of the larva from Tasmania, and a fragment of the mitochondrial *COX1* gene was amplified with primers modified from Lco1490 (5’-GGTCAACAAATCATAAAGATATTG-3’) and Hco2198 (5’-TAAACTTCAGGGTGACCAAAAAAT-3’) [10]. The amplicon was cloned with the pCR2.1-TOPO TA cloning kit (Invitrogen), the reaction transformed into chemically-competent *Escherichia coli* cells, and colonies selected on Luria-Bertani medium containing kanamycin (50 µg/ml). Plasmids were extracted from multiple *E. coli* cultures, and then the inserts sequenced using Sanger chemistry at the Australian Genome Research Facility (AGRF).

### Fungal DNA extraction and genome sequencing

Mycelium was cultured in potato dextrose broth, and genomic DNA extracted from freeze-dried mycelium. DNA from strain UoM1 was sequenced using Illumina paired reads of 125 nucleotides on a HiSeq 2500 machine by the AGRF. DNA from strain UoM4 was sequenced using Illumina paired reads of 150 nucleotides using a NovaSeq 6000 machine at the Murdoch Children’s Research Institute.

### Genome assemblies

A preliminary genome assembly of strain UoM1 was created from Illumina sequencing reads in Geneious version 8.1.9, using Velvet [11]. A *k*-mer of 71 was selected based on Velvet optimization aimed at obtaining longest contigs. Gene predictions were made using Augustus version 3.3.3 using the parameters based on *Neurospora crassa* or *Aspergillus nidulans* [12].

A second round of genome assembly was undertaken, as follows. Quality filtering and adapter removal from raw Illumina reads was conducted using BBduk (*k = 23, trimq=15, minlen=45*) from the BBmap suite v.36.86 [13]. Two approaches were adopted to produce draft genome assemblies of strains UoM1 and UoM4. First, Unicycler v.0.4.8 [14] was used, which is a SPAdes-optimiser assembler, with automatically selected *k-*mer values. A second *de novo* assembly was generated for both strains using SOAPdenovo2 v.2.40 [15] (SOAPdenovo-127mer) testing the *k-*mer length parameter between 71 and 91, in 2 bases increments. Assembled genomes were screened and filtered for potential bacterial contamination using Kraken v.2.1.1 [16]. The completeness of the genome assemblies was evaluated through identification of Benchmarking Universal Single-Copy Orthologs vis BUSCO v.1.2 [17]. Genome statistics were compared using Quast v.2.0.5 [18]. Based on the results from BUSCO analysis, genomes produced by Unicycler were selected for further analyses. OcculterCut v.1.1 [19] was used to scan the generated genomes to determine their percent GC content and distribution. The analysis was also conducted on the genome of *O. sinensis* strain IOZ07 (GenBank accession GCA_012934285.1 [20]) to compare the GC content distribution of *O. robertsii* genomes produced here.

To be able to conduct evidence-based annotation of the genomes, RNA sequencing data from *O. sinensis* was obtained from the NCBI SRA database (PRJNA687052, PRJNA673413, and PRJNA673413) and aligned against the genome of strain UoM4 using HISAT2 V.2.2.1 [21], and those reads that aligned to the genome were further extracted to produce a transcriptome assembly using Trinity v.2.10.0 [22]. Repeat libraries for both genomes were generated using RepeatModeler v.2.0.1 [23] to allow for repeat masking prior to annotation. Genome annotations were conducted using Maker v.2.31.9 [24]. The resulting annotations from the first round were further used to produce hidden Markov model (HMM) profiles for the genomes, which was further refined with a second round of SNAP training and used for the final annotation [24].

### Phylogenetic analyses

The internal transcribed spacer (ITS) regions were amplified from *O. robertsii* genomic DNA using primers ITS1 (5’-TCCGTAGGTGAACCTGCGG-3’) and ITS4 (5’-TCCTCCGCTTATTGATATGC-3’) [25], and sequenced using Sanger chemistry at the AGRF. Sequences from *Ophiocordyceps* species identified from the literature or from BLAST comparisons with *O. robertsii* sequences were downloaded from GenBank (accessions are provided in table S1). *O. agriotidis* was selected as an outgroup. *O. robertsii* strain UoM1 sequences, other than for the ITS regions, were obtained from the next generation sequencing reads. The sequences were aligned in clustalW, imported into MEGA X [26], and alignments adjusted by eye. Phylogenetic trees were created using maximum likelihood with support for relationships tested from 1000 bootstrap replicates.

For Bayesian analysis, the best nucleotide substitution model for individual loci were determined using PAUP v.4.0 [27] and MrModeltest v. 2.3 [28], to be (GTR+I+G) for *TEF, LSU*, and *RBP2*; (GTR+I) for *SSU*; and (GTR+I) for *RBP1*. Two MCMC chains were run for each individual locus and the concatenated alignment using MrBayes v. 3.2.4 [29], saving one tree per 100 generations. The run ended automatically when the standard deviation of split frequencies reached below 0.01. The 50 % majority rule consensus trees were generated after 25 % burn-in of the saved trees.

### Targeted sequence assembly and analysis of the mating type (*MAT*) locus

Defining the mating type (*MAT*) locus sequences used a combination of PCR and Sanger chemistry sequencing and next generation read assembly. First, inverse PCR was used to establish the presence of a second mating type in strains of *O. robertsii* after the identification of the putative *MAT* locus in strain UoM1. Genomic DNA of strain UoM4 was digested with restriction enzymes (EcoRI, KpnI and NcoI) then circularized using T4 DNA ligase, and this template used for PCR amplification with primers MAI0553 (5’-GCCAGAGGACTCTGCAGG-3’) and MAI0562 (5’-CCAGTGGTGGTTGTGCAGAC-3’). This established a region of DNA in strain UoM4 that was not present in the genome assembly of strain UoM1, and became a starting point for focused PCRs. Second, the genome of UoM4 was sequenced, and a targeted assembly generated based on partial sequences already obtained from PCR and sequencing and by mapping to strain UoM1. Third, the genes adjacent to the *MAT* locus and within each idiomorph were predicted using BLAST comparisons to GenBank.

Once the *MAT* locus idiomorph sequences were resolved, to test the proportion of mating types in the 11 strains derived from the sample from Tasmania in 2018 and the four strains derived from the sample in Victoria in 2019, a region of the *MAT1-1-1* gene was amplified with primers MAI0663 (5’-CTTCTGAACTTCCTCACG-3’) and MAI0664 (5’-GCAAAGTGAAAGTCGTCC-3’) and from the *MAT1-2-1* gene with primers MAI0665 (5’-AATCCCAATCCTCAGTGG-3’) and MAI0666 (5’-CGCAGTGTTGGAATCACC-3’). Amplicons were resolved on 1% agarose gels stained with ethidium bromide.

## RESULTS AND DISCUSSION

### Isolation of *Ophiocordyceps robertsii* strains into culture

Ascospores from an *O. robertsii* fruiting body collected from Tasmania in 2018 were plated onto PDA medium, and 11 strains isolated. The strains grew slowly in culture, reaching a colony diameter of about 2 cm after a month (Fig. 1A). Cultures also produced fruiting bodies, but in a stochastic and irregular manner (Fig. 1B). A number of attempts and growth conditions were used to optimize the induction of this fruiting process. The most reliable conditions observed, to date, are eight months incubation in a starting volume of 25 ml potato dextrose broth in a 20 cm high petri dish, in darkness at 14°C. All 11 strains were able to produce these structures, but they did not progress further towards the generation of sexual perithecia. *O. xuefengensis*, a heterothallic close relative of *O. robertsii*, also makes asexual stromata; the formation of these structures also have a preference for darkness [30].

**Fig 1.**
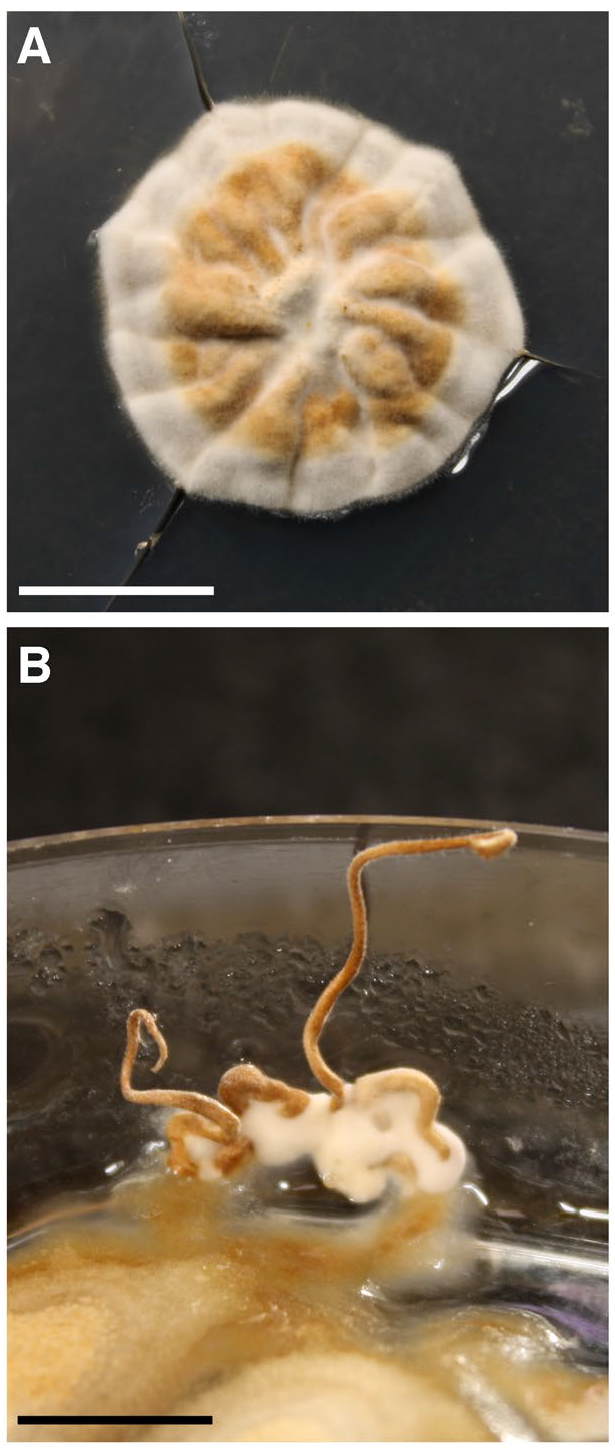
Properties of *O. robertsii* in culture. **A**. Radial vegetative growth on PDA after 32 days. Bar = 1 cm. **B**. Production of asexual stromata after ∼ 3 months. Bar = 1 cm.

To ensure the growth of these isolates was representative of the species, a second fruiting body collected in Victoria in 2019 was used to isolate four additional strains. These four strains also grew poorly, and this trait in strains from both sources is consistent with recent observations of the slow growth rate of *O. robertsii* strains isolated from New Zealand [31].

The sample from Tasmania included the parasitized insect larva, and its identification was sought by amplification, cloning and sequencing a diagnostic fragment of the *COX1* gene (GenBank accession OP376821). Comparison with the GenBank nr database revealed >99% identity to species in the ghost moth genus *Oxycanus*. The species is unknown, due to ambiguities in species identification from which DNA samples were sequenced [32, 33].

### Draft genome assemblies of *O. robertsii* indicate a larger size compared to other *Ophiocordycipitaceae* species

The genomic DNA samples of two *O. robertsii* strains (UoM1 and UoM4) were sequencing using short-read Illumina technology (deposited as GenBank PRJNA630494). Strain UoM1 was sequenced first, and a draft genome obtained after a Velvet *k*-mer optimization process designed to optimize maximum contig length. Using a *k*-mer option of 71 and a minimum cut off contig length of 1 kb, 12,910 contigs were assembled for a total genome size of 97.8 Mb. Analysis of this assembly using Augustus predicted 10,818 or 11,280 genes (*N. crassa* and *A. nidulans* parameters, respectively).

Genome assemblies constructed using Unicycler were found to have no gaps (represented by N’s in the assemblies; table S2) and resulted in more complete genomes compared to those generated with SOAPdenovo2, based on sordariomycetes_odb10 BUSCO results (97% completeness for UoM1 and 97.2% completeness for UoM4) (table S3), and therefore were selected for subsequent genome annotation and OcculterCut analyses. The differences in genome predictions likely reflect the differences in the underlying assembly parameters. Unicycler was originally developed for bacterial genome assembly [14], hence, may outperform SOAPdenovo in haploid genome assembly. Unicycler assembly for strain UoM1 consisted of 10,216 contigs and a total size of 95.8 Mb, while the assembly for strain UoM4 included 5,720 contigs with a total genome size of 102.7 Mb. These assemblies are in GenBank as BioProject PRJNA630494 (JAPEBV000000000 for UoM1 and JAPEBW000000000 for UoM4). Analyses of the assemblies using evidenced-based annotation in Maker predicted 8,904 and 8,934 genes for UoM1 and UoM4, respectively.

The distribution of genes within the *O. robertsii* genome assemblies is uneven, with many of the largest contigs having no genes predicted to be present on them, and a pattern that is consistent with previous observations of large genome expansions in the Ascomycota not correlating with increased gene content [34]. This correlates with AT content, with contigs being unevenly distributed in AT vs. GC content, and correspondingly genes not predicted in the AT rich regions (Fig. 2). OcculterCut results showed that the GC content distribution in both genomes was bimodal, similar to that of *O. sinensis* strain IOZ07 (Fig. 2A, B, C). However, the proportion of AT-rich regions in *O. sinensis* strain IOZ07 (78%) was higher compared to that of UoM1 (70%) and UoM4 (72%). OcculterCut analysis also found gene density in the AT-rich regions of UoM1 and UoM4 was 0.148 and 0.081 genes per Mb, respectively, while gene density in the GC-balanced regions of these strains was 312 and 313 genes per Mb (example shown in Fig. 2D). This ‘bipartite’ structure is reminiscent of the genomes of a number of fungi, such as *Leptosphaeria maculans*, a plant pathogen in which this genome structure was first observed [35]. The structure is attributed to repetitive regions, usually transposable elements, being targeted by repeat induced point (RIP) mutation, converting cytosine to thymine and effectively raising the AT content. RIP mutation is predicted to occur in many fungi, and has been experimentally demonstrated in the classes Sordariomycetes, Dothideomycetes and Eurotiomycetes [36-38]. RIP occurs in *Trichoderma reesei* [39], which is in the same order (Hypocreales) as *O. robertsii*.

**Fig 2.**
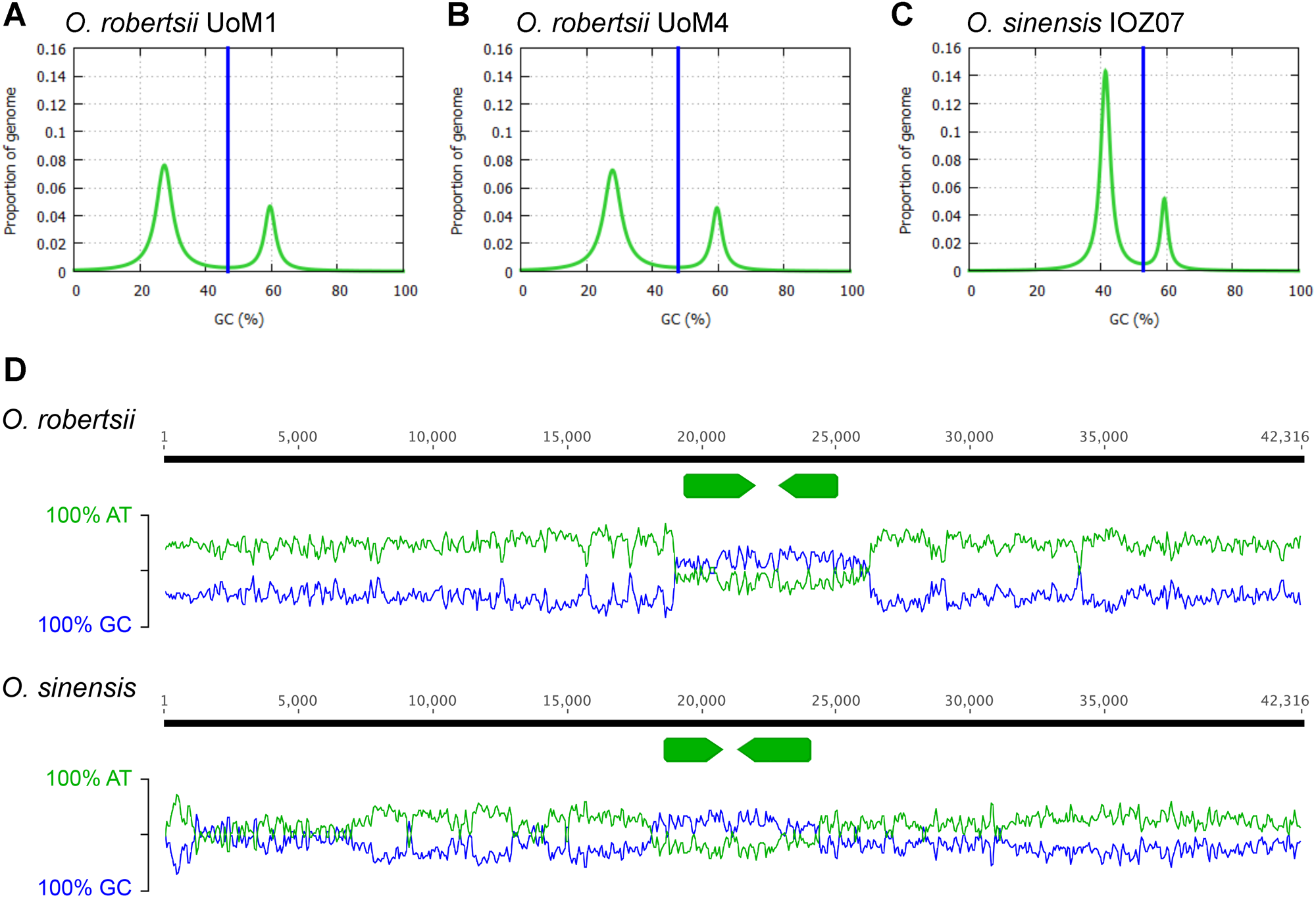
The GC content distribution of *O. robertsii* strains in bimodal. **A**. Strain UoM1 and **B**. strain UoM4 showing a similar structure to that of **C**. *O. sinensis* strain IOZ07. Vertical blue lines show the GC cut-off points selected by OcculterCut [19] to classify genome segments into distinct AT-rich and GC-balanced regions. **D**. An example of an *O. robertsii* contig that illustrates its bipartite genome, comprising AT-rich and gene poor regions vs. GC-rich and gene dense regions. AT and GC content is measured using 100 bp sliding windows, with two genes marked with the arrows. The homologous region from *O. sinensis* strain IOZ07 is shown for comparison; note that the AT-rich regions extend for more than 30 kb and 80 kb on either side of the two genes. BLASTx of the surrounding region returns hits in GenBank to retrotransposable elements; in the two species they feature numerous stop codons.

The genome sequence of *O. robertsii* appears to be considerably larger than other species in the *Ophiocordycipitaceae* family, which typically range between 21.9 to 49.3 Mb [40, 41], but is similar to that of the far larger *O. sinensis*. There have now been at least five genome sequencing attempts on *O. sinensis* towards complete coverage, with the best current estimate of a genome sized at 110-120 Mb [9, 20, 42-44]. As a consequence of the well-established challenge to sequence *O. sinensis*, attention was not placed on trying to optimize further genome assembly parameters for the Illumina short reads of *O. robertsii*, as a better understanding of the genome structure, genome expansion, gene content, and gene arrangements relative to *O. sinensis* will require generation of a high quality version of the genome by incorporating long read sequencing technologies coupled to RNA sequencing to aid in gene annotation. However, some points are of relevance in comparing *O. robertsii* and *O. sinensis*. One is that more than two thirds of the *O. sinensis* genome is comprised of repetitive elements, and the species is proposed to have an active RIP mutation system, and, unlike in other fungal species, is hypothesized to have targeted the DNA repeats for the ribosomal RNA molecules for RIP, which has led to difficulties using the ITS regions for phylogenetic studies [42, 45, 46].

It has been hypothesized that the expansion in genome size in *O. sinensis* occurred during the rising in altitude of the Tibetan Plateau, and that the genome expansion contributed to its adaptation to this environment [9, 42]. However, *O. robertsii* has a similar genome structure, and one set of the strains were isolated from Queenstown, which is 129 m above sea level. Furthermore, there are just under 500 reports of *O. robertsii* in the Atlas of Living Australia (www.ala.org.au), a citizen-scientist database, all from southeastern Australia and New Zealand locations well below the altitude of the Tibetan Plateau.

### Phylogenetic analyses place *O. robertsii* as one of the close relatives of the medicinal fungus *O. sinensis*

Sequences of six DNA regions were obtained from GenBank for close relatives of *O. robertsii*, such as *O. sinensis, O. xuefengensis* [47], *O. macroacicularis* [48], *O. lanpingensis* [49], *O. karstii* [50] and others, and then compared with those same regions from *O. robertsii* by alignment and the construction of phylogenetic trees.

The six trees show limited concordance between them for the relationships between the species and often low bootstrap support or Bayesian posterior probabilities (Fig. S1), which has been observed previously [see for example the phylogeny of *Ophiocordyceps* by [51]]. Concatenation of five of the regions (i.e. excluding the ITS region) yielded a tree with increased support at many nodes (Fig. 3). Different sets of genes, or potentially whole genome comparisons, are required if using DNA sequencing information to provide stronger relationship indicators between *Ophiocordyceps* species.

**Fig 3.**
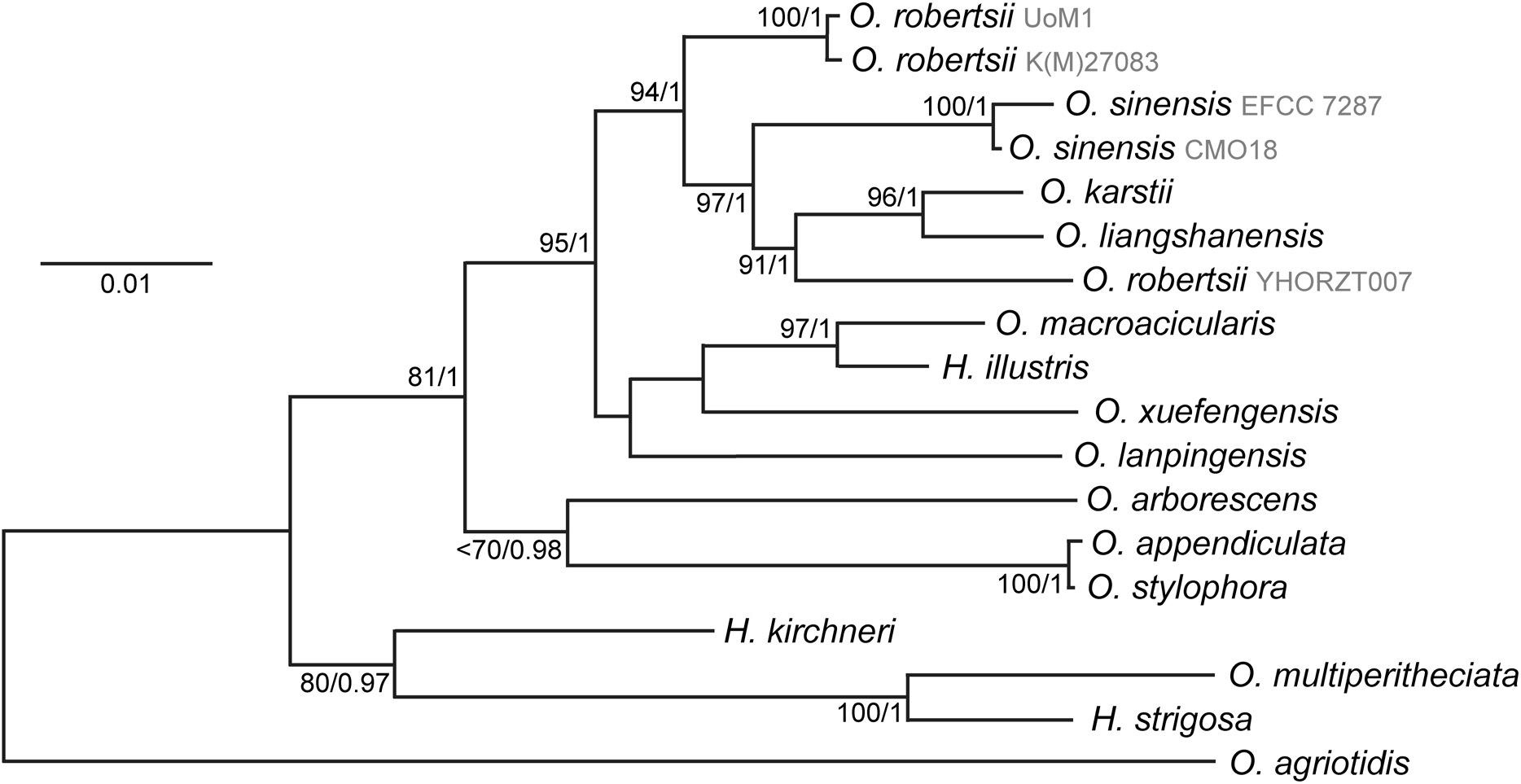
Phylogenetic tree of the relationships between *Ophiocordyceps* species closely related to *O. robertsii* and *O. sinensis* generated using maximum likelihood. *H* is the abbreviation for *Hirsutella. O. agriotidis* was used as the outgroup. Concatenation of DNA sequences of regions of *SSU, LSU, TEF1α, RBP1* and *RBP2*. Numbers adjacent to nodes indicate bootstrap % from 1000 replicates if above 70% / Bayesian posterior probability if above 0.95. The General Time Reversal +G model was used.

Several points emerge from the phylogenetic analyses. The first of these is that *O. robertsii* is not the closest relative to *O. sinensis*, at least based on the phylogenetic trees generated. The strains isolated in this work have identical ITS sequences (GenBank accession for the strains derived from Victoria is OP326271) and are very similar to *O. robertsii* collected in 1994 from Australia, while YHORZT007 from China [49] is misidentified as *O. robertsii* and instead may represent a novel species. The ITS sequences in GenBank from *O. robertsii* strains from New Zealand share less than 97% identity with Australian strains, which is potentially an indication that specimens classified as *O. robertsii* may be two distinct species in the two countries. Another *Ophiocordyceps* species that is entomopathogenic and endemic to New Zealand is *Cordyceps hauturu* [52]. While similar to *O. robertsii* in appearance, the one ITS sequence available is only 91% similar to that of *O. robertsii* and therefore is a distinct species. One final point is that *O. stylophora* and *O. appendiculata* are likely the same species.

### Analysis of the predicted mating type (*MAT*) locus of *O. robertsii* supports a heterothallic reproductive strategy

The genome sequencing assembly of strain UoM1 was searched, using BLAST, for homologs of the mating type genes or two flanking genes that are adjacent to the *MAT* locus in ascomycetes. One region was identified, and initial PCR analysis suggested that this was found in only a subset of the *O. robertsii* strains. To obtain additional *MAT* sequences, inverse PCR was used to identify the idiomorphic region in strain UoM4. Due to challenges and limited efficiency in this method, strain UoM4 was then also sequenced and assembled, with attention paid to resolving the *MAT* region by a combination of reiterative targeted assemblies of short reads coupled to PCRs and sequencing of those amplicons. The DNA sequence was completed except for one region that could be amplified by PCR, but could not be fully resolved by sequencing off those amplicons, in one of the strains (this gap is marked in Fig. 4A). The sequences of the two *MAT* loci have been deposited to GenBank as accessions MT436030 and MT436031.

**Fig 4.**
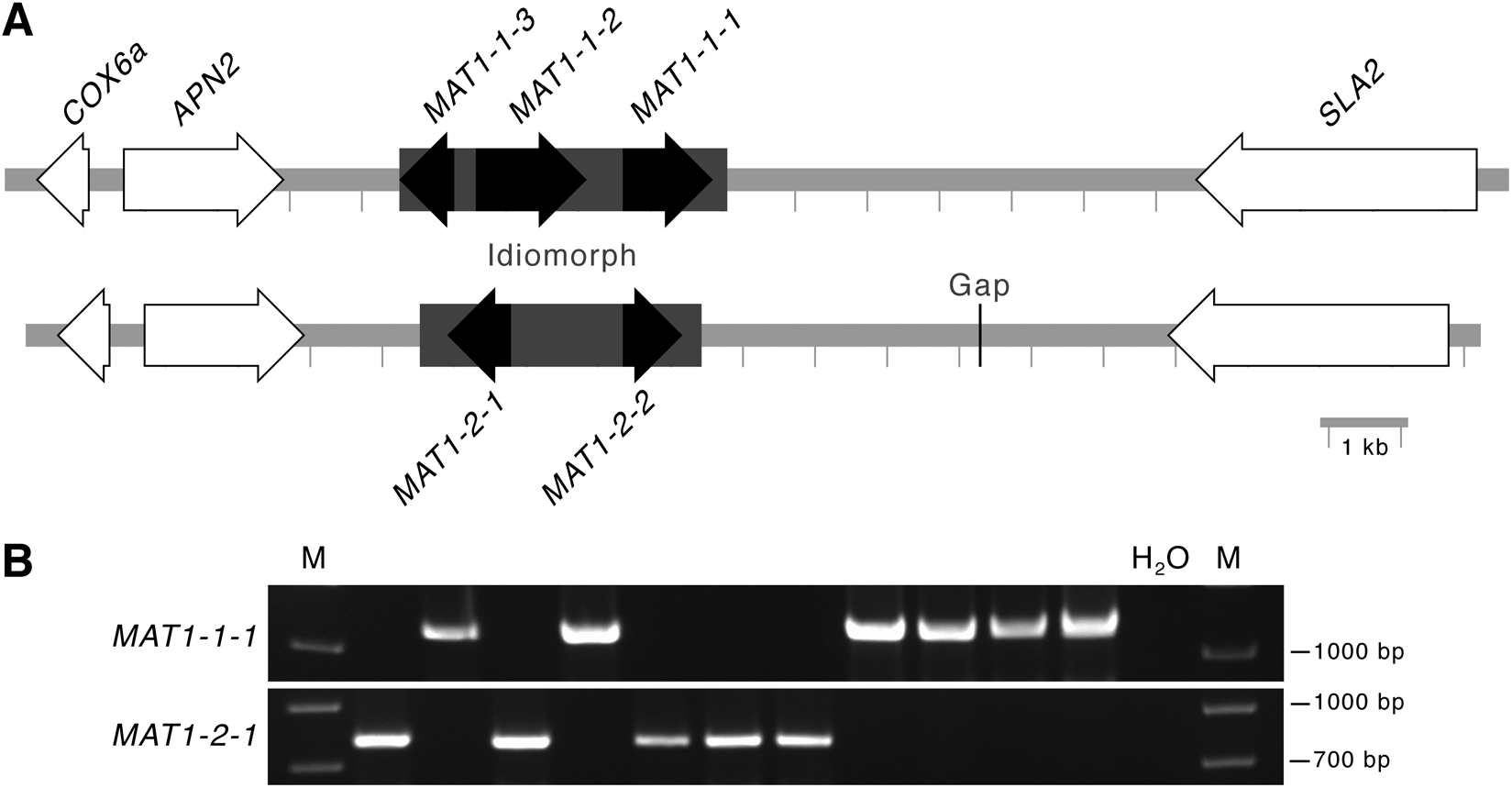
DNA sequence analysis suggests that *O. robertsii* has a heterothallic life cycle. **A**. Diagram of the *MAT* locus from two strains of *O. robertsii*. The genes in white indicate those found adjacent to *MAT* loci in other fungi. DNA sequence similarity is low in the idiomorph region (grey box), in which one strain encodes three predicted genes and other encodes two predicted genes. The gap indicates a region of low sequence coverage. **B**. Agarose gels resolving PCR amplification products using primers specific to either *MAT1-1-1* or *MAT1-2-1* from 11 isolates of *O. robertsii*. M indicates the DNA ladder; H_2_O indicates no DNA was added to the PCRs.

Comparison of the two *O. robertsii* strains indicates that each contains an idiomorphic DNA region that has minimal similarity between each other (Fig. 4A). The *MAT1-1* idiomorphic region of 4.5 kb contains three predicted genes similar to those found in *MAT* loci of other fungi. The *MAT1-2* idiomorphic region of 3.9 kb contains two predicted genes. Comparison of the predicted proteins within the *MAT* locus by BLAST against GenBank returned highest matches to *O. sinensis* and *O. xuefengensis*, which is consistent with the inter-species relationships observed using other genes (Fig. 3).

*MAT1-1-1* encodes the ‘MATα’ high mobility group (HMG)-box protein that is a fundamental determinant of mating type (e.g. as α in *Saccharomyces cerevisiae*). *MAT1-1-2* is commonly found in *MAT* loci in the Sordariomycetes: deletion of the homolog in *Cordyceps militaris* results in a strain able to form fruiting bodies but with sterile perithecia [53]. *MAT1-1-3* encodes another HMG-domain protein (MATA-type).

The *MAT1-2-1* gene encodes the HMG-domain protein (MATA-type) found in ascomycete *MAT* loci. The *MAT1-2-2* gene encodes a protein of unknown function that is not commonly associated with *MAT* loci in ascomycete species. The sequence available for the *MAT1-2* idiomorph of *O. xuefengensis* is likely truncated [54], so it is unclear if the homolog is present in this species. However, BLAST of genome sequences available for other *Ophiocordycep*s species and analysis of the flanking genes indicates that *MAT1-2-2* is also found in the predicted *MAT* locus of *O. sinensis* (at least in strain CO18), *O. australis* and *O. camponoti-rufipedis*.

Fragments of the *MAT* regions were amplified from the 11 strains isolated from the fruiting body collected in Tasmania in 2018. Each contained only one of the regions (Fig. 4B). Of the four strains isolated from the fruiting body from Victoria in 2019, two were *MAT1-1* and two were *MAT1-2* (data not shown). Hence, both the DNA sequence information and idiomorph distribution are consistent with *O. robertsii* having a heterothallic lifestyle.

During our research, the nature of the *MAT* locus of *O. xuefengensis* was explored indicating that this species is also heterothallic [54, 55]. The sexual reproduction system in *O. xuefengensis* has been characterized, and sheds light on the fruiting structures formed by single strains of *O. robertsii*. That is, *O. xuefengensis* and presumably *O. robertsii* produce an asexual stroma that is subsequently fertilized by the opposite mating type. With slow growth in culture, completion of the *O. robertsii* life cycle *in vitro* is yet to be achieved. The resources developed here, including genome assemblies that are available from GenBank and strains, one of each mating type that have been deposited in the Queensland Herbarium culture collection (BRIP 75182 a and BRIP 75183 a), will contribute to this endeavor.

## Conclusions

*Ophiocordyces robertsii* is endemic to Australia and New Zealand, and a notable species in New Zealand’s culture. While this research focused on cultivation of the species, for the prospective benefits such cultures could bring, it also has ramifications about the evolution and biology of *O. sinensis*, which is one of the world’s most expensive fungi. *O. sinensis* has been difficult to cultivate *ex situ* due to its niche geographical range, and only a handful of companies have commercial production [5, 6]. This research established that *O. robertsii* could serve for comparative analysis with *O. sinensis* and other *Ophiocordyceps* species. The converse is that the medicinal benefits associated with *O. sinensis* are underexplored in *O. robertsii*. Last, it is currently unclear if *O. robertsii* in Australia and New Zealand represents one or multiple species.

## ACKNOWLEGMENTS

We thank Christine Wilson for finding samples of *O. robertsii*.

---

**Fig S1.**
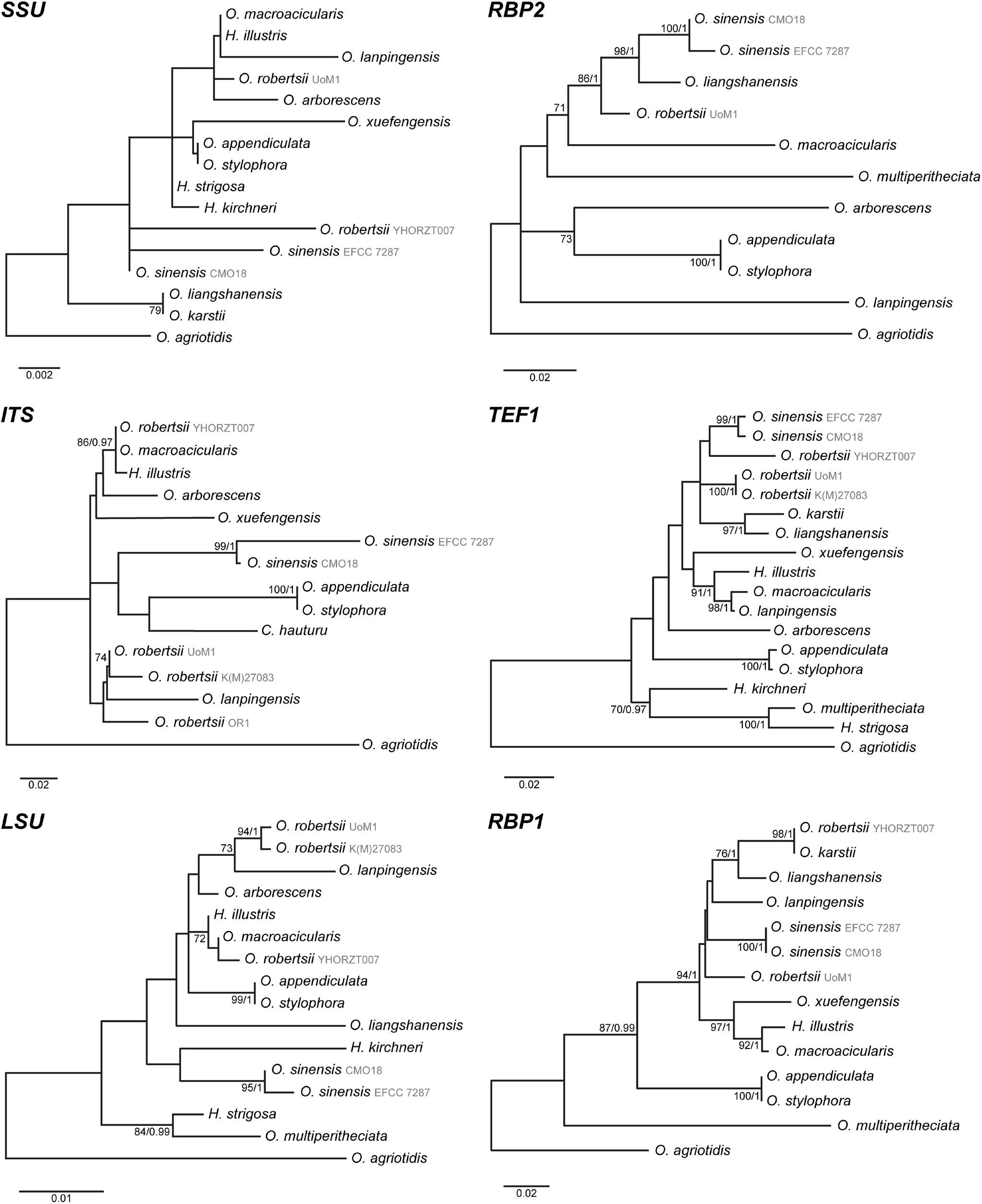
Phylogenetic trees of the relationships between *Ophiocordyceps* species closely related to *O. robertsii* and *O. sinensis* generated using maximum likelihood. *H* is the abbreviation for *Hirsutella* and in the ITS tree *C* is the abbreviation for *Cordyceps. O. agriotidis* was used as the outgroup. Numbers adjacent to nodes indicate bootstrap % from 1000 replicates if above 70% / Bayesian posterior probability if above 0.95. Models used were Kimura-2 +G (*SSU*), Tamura-Nei +G (*RBP2, TEF1*), Hasegawa-Kishino-Yano +G (*ITS, LSU*) and Tamura-3 (*RBP1*).

